# Oral delivery of GLP-1R agonist by an engineered probiotic yeast strain has anti-obesity effects in mice

**DOI:** 10.1101/2022.12.21.521368

**Authors:** Karl Alex Hedin, Hongbin Zhang, Vibeke Kruse, Vanessa Emily Rees, Fredrik Bäckhed, Thomas U. Greiner, Ruben Vazquez-Uribe, Morten Otto Alexander Sommer

**Affiliations:** Novo Nordisk Foundation Center for Biosustainability, Technical University of Denmark, 2800, Kgs. Lyngby, Denmark; The Wallenberg Laboratory, Department of Molecular and Clinical Medicine, Institute of Medicine, Sahlgrenska Academy, University of Gothenburg, Gothenburg, Sweden; Region Västra Götaland, Sahlgrenska University Hospital, Department of Clinical Physiology, Gothenburg, Sweden; Novo Nordisk Foundation Center for Basic Metabolic Research, Faculty of Health Sciences, University of Copenhagen, Copenhagen, Denmark

## Abstract

Obesity is rapidly increasing within the global population and is one of the leading causes of chronic diseases, including type 2 diabetes (T2D), non-alcoholic fatty liver disease, and cardiovascular diseases. Glucagon-like peptide-1 receptor (GLP-1R) agonists have emerged as promising therapeutic agents for treating T2D and obesity. However, the route of administration of the GLP-1R agonists is currently by injection or high oral dosages of the therapeutic combined with absorption enhancers. Oral delivery of GLP-1R agonists remains the preferred administration route due to convenience and high patient compliance. Thus, strategies to improve the oral delivery of this therapeutic are needed. In this study, we engineered the probiotic yeast *Saccharomyces boulardii* strain to produce Exendin-4, a GLP-1R agonist, in the gastrointestinal tract to reduce the adverse effects of diet-induced obesity in male C57BL/6 mice. The biological efficiency of the secreted Exendin-4 from *S. boulardii* was characterised *ex vivo* on isolated pancreatic islets, demonstrating induced insulin secretion. Furthermore, *in vivo* characterisation of the engineered strain identified a synergistic effect of cold exposure and Sb-Exe4 by successfully inhibiting appetite and promoting body weight loss under cold exposure (8°C). In addition, the combination of cold and Sb-Exe4 improved the glucose and lipid homeostasis in the mice by increasing the circulating glucagon level and reducing the inflammatory marker TNF-α. Our results demonstrate that *S. boulardii* can be genetically modified to secrete and deliver active therapeutic GLP-1R agonists in the gastrointestinal tract improving the metabolism of the host.

## Introduction

Obesity has become one of the top public health concerns worldwide, affecting more than 13% of the population globally^1,2^. Obesity is defined as excessive fat accumulation and is one of the major contributors to chronic diseases, including type 2 diabetes (T2D), non-alcoholic fatty liver disease, and cardiovascular diseases^3–6^. Diet and exercise are considered the foundation for the prevention of obesity; however, most patients do not experience sustained results with these interventions^7^. In addition, these interventions have been insufficient in some patients due to their genetics^8^ or microbiota^9^, which might directly affect the severity of the disease. Furthermore, surgical procedures and alternative therapies have been deployed over the past decades to treat obesity^10^. However, the majority of alternative therapies have been withdrawn from the market due to substantial adverse effects^10,11^. Glucagon-like peptide-1 (GLP-1), which is produced in the intestinal epithelial endocrine L-cells and secreted in response to ingestion of nutrients^12^, has shown great promise in the treatment of obesity as well as other diseases^13,14^. However, the short half-life (∼2 minutes) of GLP-1 in the blood has limited its clinical value for T2D and obesity treatment^12,15^. Past efforts have focused on developing GLP-1 receptor (GLP-1R) agonists with improved half-life^16^, including alternatives such as the long-acting potent GLP-1R agonist, Exendin-4, with an estimated half-life of 90 –216 min^17^. Nonetheless, the route of administration of Exendin-4 is via twice-daily subcutaneous injection^18^; which is a less preferred administration route with lower adherence among patients^19,20^. Although oral administration has been extensively studied^21–24^, cost efficiency and bioavailability remain challenging^25^. Thus, new oral delivery strategies that can achieve stable and therapeutically relevant systemic exposure levels *in vivo* are of great therapeutic interest.

Advanced microbiome therapeutics (AMTs) is an emerging field of medicinal synthetic biology which utilises engineered microbes to perform therapeutic functions on and within the human body^26^. AMTs have previously been engineered for the delivery of peptides in the gastrointestinal tract of rodents; for example, *Lactococcus lactis* and *Lactobacillus gasseri* have been engineered to secrete GLP-1^27–29^, and *Escherichia coli* Nissle 1917 have been engineered to secrete N-phosphatidylethanolamine^30^ and a GLP-1 analogue^31^.

In this study, we explore *Saccharomyces boulardii* as potential AMT chassis for the oral delivery of therapeutic peptides. *S. boulardii* is a probiotic yeast with favourable properties to ameliorate certain diseases associated with phenotypes observed in *db/db* mice^32^. Its close relative, *Saccharomyces cerevisiae*, is one of the most used hosts for biopharmaceutical synthesis, gene manipulation and protein production^33,34^. Since recombinant tools developed for *S. cerevisiae* can be applied in *S. boulardii*^35–37^, the organism is well suited as an AMT chassis. *S. boulardii* has previously been engineered to secrete IL-10 and atrial natriuretic peptides to enhance its anti-inflammatory properties^38^. Here, we sought to engineer *S. boulardii* to deliver a GLP-1R agonist in the gastrointestinal tract of mice fed a high-fat diet, as a proof of concept to investigate the potential anti-obesity effects of a genetically modified *S. boulardii*.

## Results

### *S. boulardii* secretes high levels of biologically active GLP-1R agonists *in vitro*

We started by engineering *S. boulardii* to produce and secrete GLP-1R agonists (Figure 1A). To promote secretion of the peptides, we fused the genes to the mating alpha-factor secretion leader^39^ under a strong constitutive *TDH3* promoter^40^ and the highly active *DIT1*\*\* terminator^41^. We inserted a KOZAK sequence before the alpha-factor secretion leader to increase the translational efficiency^42^, a Kex2 site^43^ and a spacer peptide^44^ downstream of the alpha-factor secretion leader to increase cleavage efficiency^45^. In addition, we evaluated the effect of multiple chromosomal integrations of the expression cassette and compared single, double and quadruple integrations. The number of chromosomal copies of the gene increased the concentration for both GLP-1 and Exendin-4 (Supplementary Figure S1A; D). However, multiple copies also resulted in a higher metabolic burden for the strains (Supplementary Figure S1B; E), resulting in a lower growth rate (Supplementary Figure S1C; F). Therefore, we chose to proceed with the double integration strain to balance the growth rate and product secretion to obtain a robust strain^46^. These engineered double-integrated strains produced 16 nM/OD_600_ of GLP-1 and 15 nM/OD_600_ of Exendin-4 in aerobic fermentation and 5 nM/OD_600_ of GLP-1 and 11 nM/OD_600_ of Exendin-4 in anaerobic fermentation (Figure 1B). The *S. boulardii* strain containing the empty vector (Sb-Empty) displayed no levels of GLP-1 or Exendin-4.

**Figure 1.**
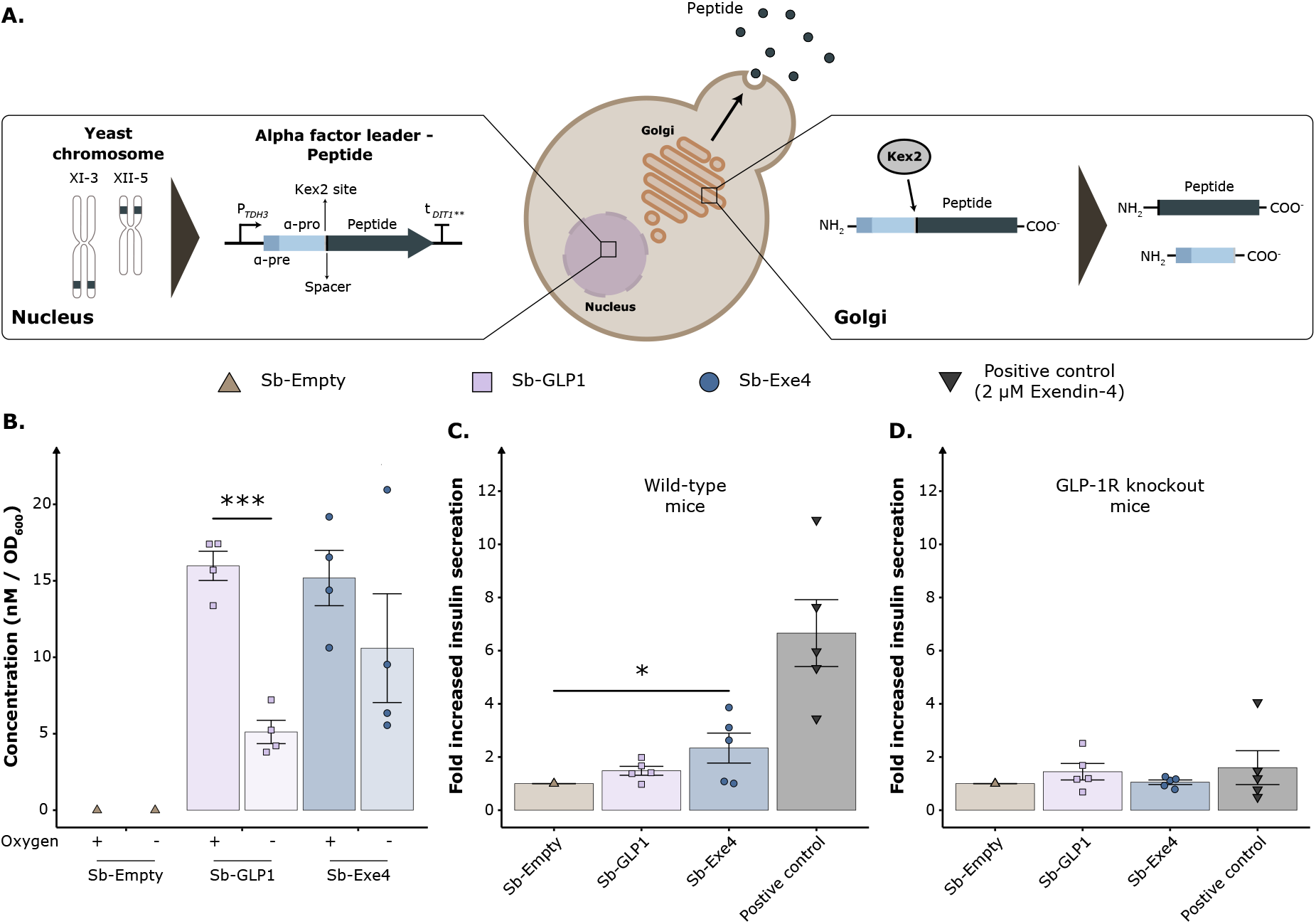
*In vitro* and *ex vivo* characterisation of the *S. boulardii* secreting GLP-1R agonists. **(A)** Schematic overview of the engineered S. *boulardii* for production and secretion of peptides. **(B)** The concentration of GLP-1 (n = 4) and Exendin-4 (n = 4) secreted by *S. boulardii* in aerobic and anaerobic conditions. **(C)** The fold change of insulin secreted normalised to the Sb-Empty, from isolated pancreatic islets of wild-type C57BL/6J mice (n = 5), **(D)** and GLP-1R knockout C57BL/6J mice (n = 5). 2 µM recombinant Exendin-4 was used as a positive control. All stimulations were performed in duplicates. Bar plots are displayed as the mean ± SEM. * p < 0.05. B was analysed with dependent sample t-test with Bonferroni adjustment for multiple comparisons. C and D were analysed with one-way ANOVA with Dunnett’s post hoc test, with Sb-Empty as a reference.

As we confirmed that the GLP-1R agonists were produced and secreted by the *S. boulardii* strains, we next sought to validate the biological efficiency of the *S. boulardii* produced GLP-1R agonists. Primary pancreatic islets isolated from wild-type mice and GLP-1R knockout mice were stimulated with supernatant from the different producer strains and compared against Sb-Empty strain as negative control and 2 µM of recombinant Exendin-4 as a positive control. Primary pancreatic islets, isolated from the wild-type mice, showed significantly higher insulin secretion after stimulation with supernatant from *S. boulardii* secreting Exendin-4 (Sb-Exe4) compared to Sb-Empty. *S. boulardii* secreting GLP-1 (Sb-GLP1) showed no effect. Similar stimulation was not observed in the islets isolated from GLP-1R knockout mice for either Sb-GLP1, Sb-Exe4, or recombinant Exe-4 (Figure 1D).

### Sb-Exe4 reduces food intake and induces weight loss during cold exposure

We sought to evaluate the therapeutic potential of Sb-Exe4 strain in the gastrointestinal tract of mice fed a high fat diet. Our previous work has established that *S. boulardii* has a short residence time in the gastrointestinal tract of a mouse, reducing in four orders of magnitude of viable cells within 24 hours^47^. Therefore, to maintain a significant effect of the Sb-Exe4 in mice, we sought to boost the number of viable cells during the mice’s most active phase^48^. We conducted a pilot study to determine the dosing regimen for *S. boulardii*. We determined that daily dosing of *S. boulardii* in the afternoon would be appropriate, as we observed a one-order-of-magnitude decrease of *S. boulardii* in the faeces during the first half of the day, followed by three orders of magnitude washout during the second half of the day (Supplementary Figure S2).

Next, we attempted to assess the effects of Sb-Exe4 daily dose over a period of 35 days in mice fed a high-fat diet, however, we did not observe any anti-obesity effect of the Sb-Exe4 treatment compared to Sb-Empty (Supplementary Figure S3). Therefore, we sought to investigate if a change in the environmental temperature might enhance the effect of Sb-Exe4 in mice. It is known that the bioenergetic profiles of animals (such as body weight, food intake, and oxygen consumption) are tightly modulated by environmental temperature^49–51^. For instance, in response to cold exposure, food intake increases in mice to compensate for the induced energy expenditure to maintain normal body temperature^49^. It has previously been demonstrated that GLP-1R agonists reduce food intake via intestinofugal neurons^52^. Therefore, we hypothesise that the produced Exendin-4 in the gastrointestinal tract activates GLP-1R and prevents the increased food intake caused by cold exposure. The mice were housed at room temperature for 24 days, fed a high-fat diet and gavaged daily with Sb-Exe4 or Sb-Empty. The mice were exposed to cold temperature (8°C; Figure 2A) 5 days before termination. The combination of cold exposure and Sb-Exe4 treatment resulted in a 4-fold higher body weight loss (Figure 3B, C) and a 25% reduced food intake (Figure 3D, E). In addition, receiving the Sb-Exe4 decreased the respiratory exchange ratio (RER; VCO_2_/VO_2_) compared to the Sb-Empty treated mice (Figure 3G, H), promoting an increased amount of fatty acids as an energy source relative to glucose. Moreover, the Sb-Exe4 mice exhibited a tendency for decreased energy expenditure and a significant increase in activity during cold exposure (Supplementary Figure S4).

**Figure 2.**
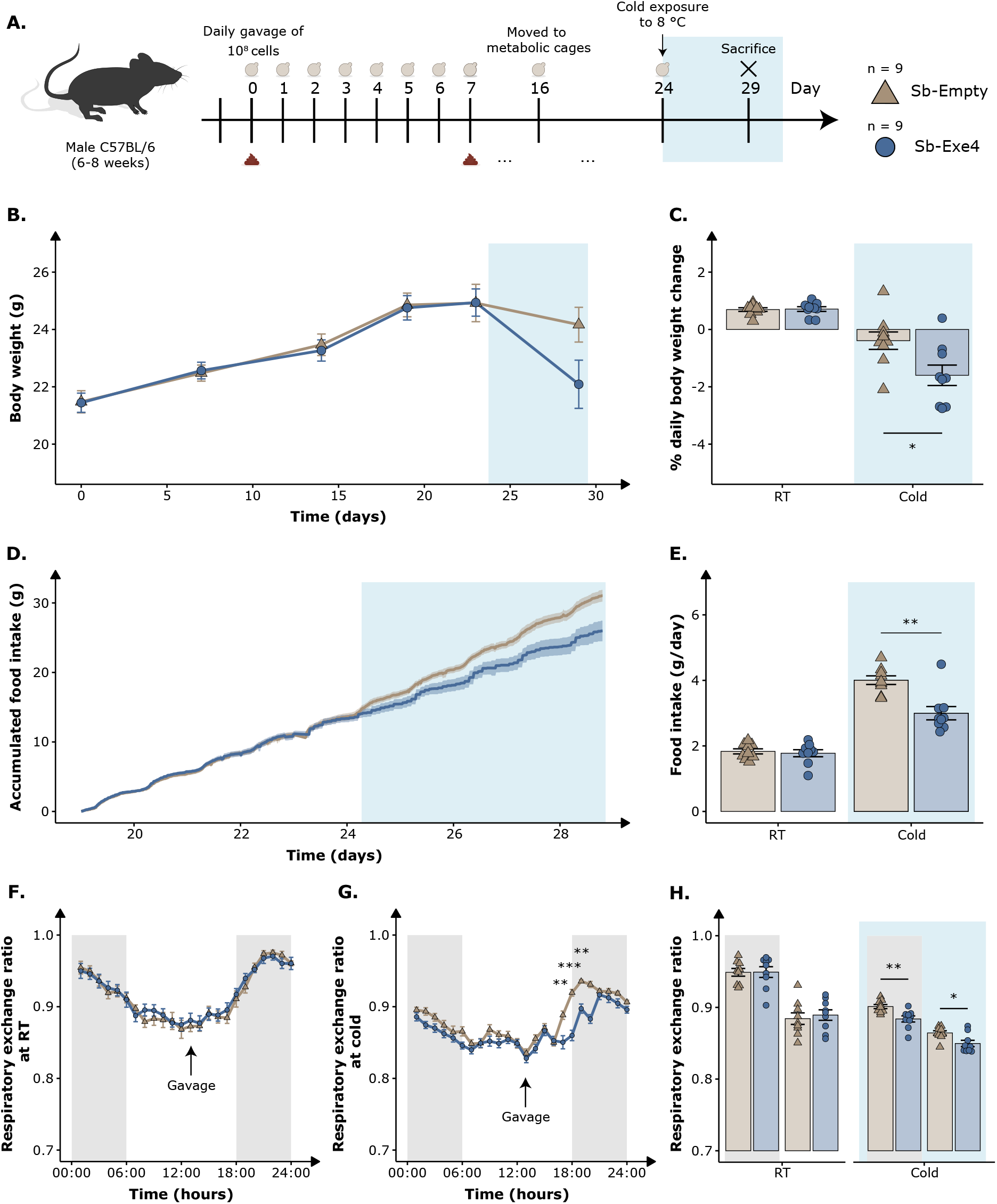
Synergetic effect of Sb-Exe4 and cold exposure on energy homeostasis. **(A)** Schematic overview of the study plan. Male C57BL/6 mice were divided into two groups, orally administered with either Sb-Empty or Sb-Exe4. Mice were initiated on a 45 %kcal high-fat diet on day 0. **(B)** Body weight was monitored weekly for 29 days while the mice were fed a high-fat diet and orally administered their respective strain. **(C)** Mean daily body weight change at room temperature (RT; 22°C) and cold (8°C). **(D)** Accumulated food intake during the recorded period. **€** Mean food intake per day at RT and 8°C. **(F)** Mean hourly respiratory exchange ratio (RER; VCO_2_ / VO_2_) under a 12:12 light-dark cycle over a 5-day period at RT and **(G)** a 5-day period at cold. **(H)** Mean RER during light and dark cycle at RT and cold. The black-shaded area indicates a dark period. The light blue shaded area indicates cold exposure (8°C). Data are presented as the mean ± SEM (n = 9). * p < 0.05, ** p < 0.01 All samples were analysed by Wilcoxon signed-rank test with Bonferroni adjustment for multiple comparisons.

**Figure 3.**
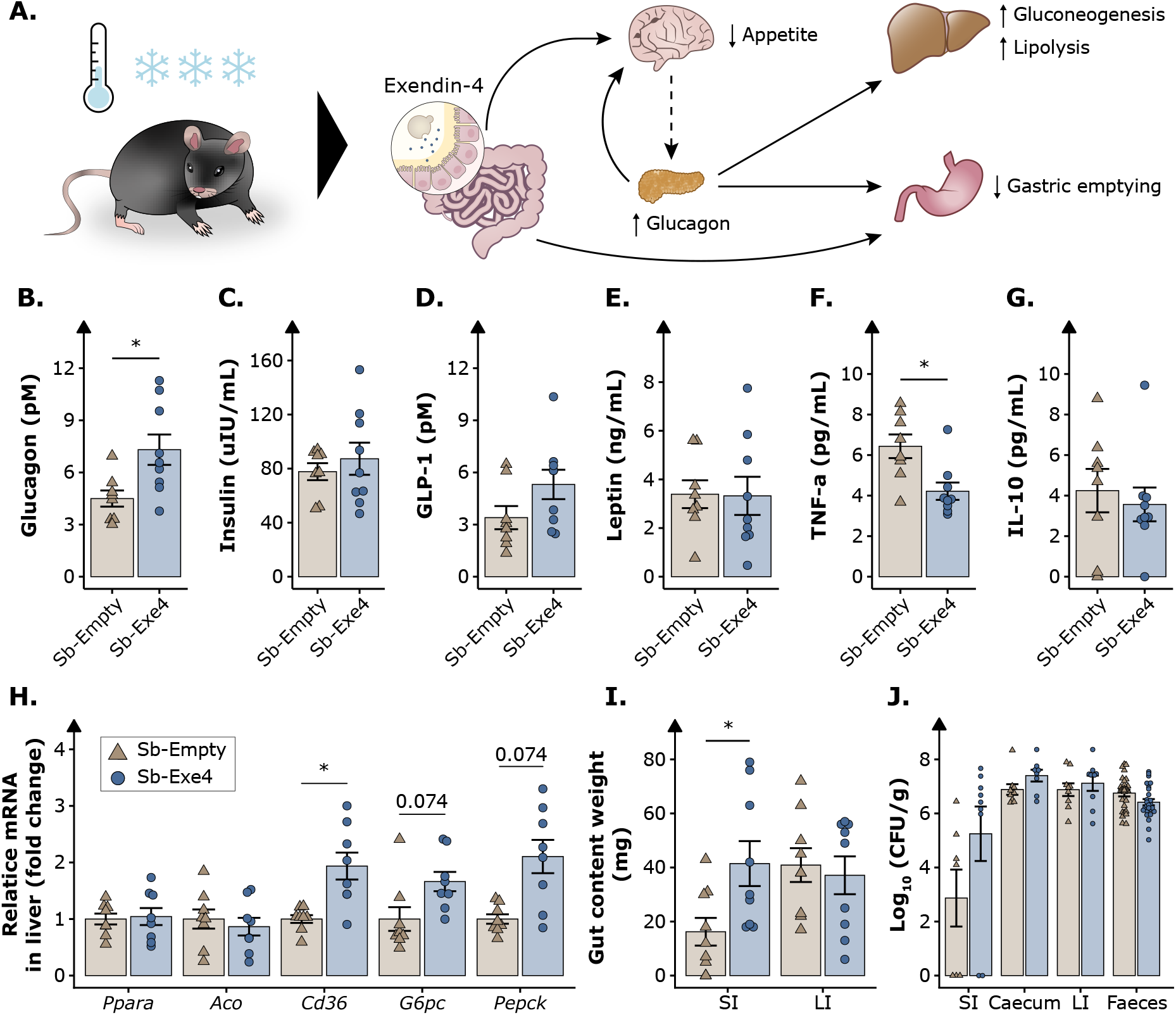
End-point characterisation of the synergistic effect of cold exposure and Sb-Exe4 in male C57BL/6 mice for 29 days. **(A)** Graphic simplification of the mechanistic effect on glucose homeostasis in male C57BL/6 mice orally administered with Sb-Exe4 and fed a 45 % kcal high-fat diet for 29 days. Fasting vena cava levels of **(B)** glucagon (pM), **(C)** insulin (uIU/mL), **(D)** GLP-1 (pM), **(E)** leptin (ng/mL), **(F)** TNF-α (pg/mL) and **(G)** IL-10 (pg/mL) at the end of the study quantified with Meso Scale Discovery (n = 8 – 9). **(H)** Relative mRNA of *Ppara, Aco, Cd36, G6pc* and *Pepck* in the liver at the end of the study. Data presented as fold change normalised to Sb-Empty (n = 9). **(I)** Weight (mg) of gut content was collected from the small intestine (SI) and large intestine (LI) of the mice (n = 9). **(J)** Abundance (Log_10_ CFU/g faeces) of Sb-Empty and Sb-Exe4 in SI (n = 9), caecum (n = 9), LI (n = 9), and faeces (n = 27) in the mice. Data are presented as the mean ± SEM. P-value written out < 0.1, * p < 0.05, ** p < 0.01. B to H and J were analysed by Wilcoxon signed-rank test, and I were analysed with dependent sample t-test. Bonferroni adjustments were used for multiple comparisons.

Furthermore, to study the long-term effect of receiving Sb-Exe4 in mice fed a high-fat diet, an additional study was conducted for 55 days in both room temperature and cold. The prolonged treatment of Sb-Exe4 does not further increase the response already observed from the shorter treatment at room temperature. However, the cold exposure in this cohort of mice showed a similarly significant effect. Mice receiving Sb-Exe4 showed a 4-fold higher body weight loss, decreases in RER, and energy expenditure (Supplementary Figure S5).

### Synergetic effect of cold exposure and Sb-Exe4 alter glucose and lipid homeostasis in mice fed a high-fat diet

To further investigate the functional association between long-term cold exposure and Sb-Exe4, we analysed the end-point biomarkers involved in energy homeostasis (Figure 3A)^53–56^. In the blood circulation, we observed a significantly increased concentration of glucagon and decreased concentration of the inflammatory cytokine marker TNF-α (tumour necrosis factor-alpha) in mice receiving Sb-Exe4 (Figure 3B-F). In line with the increased glucagon levels, the hepatic gluconeogenesis genes *G6pc* (glucose-6-phosphatase catalytic subunit) and *Pepck* (phosphoenolpyruvate carboxykinase) showed an increased tendency in expression levels. In addition, we observed an increased hepatic expression of the fatty acid transporter gene *Cd36* (cluster of differentiation 36). Mice receiving Sb-Exe4 also significantly increased the gut content mass in the small intestine. Furthermore, the prolonged 55-day treatment of Sb-Exe4 showed a similar end-point effect of cold exposure, demonstrating an increased level of glucagon, decreased level of TNF-α, hepatic induction of *G6pc* and *Cd36* (Supplementary Figure S6).

It is well established that cold exposure effectively stimulates the UCP1-mediated thermogenic energy expenditure^57^. Moreover, increased GLP-1R agonist and glucagon levels have previously been reported to increase the browning of subcutaneous white adipose tissue (sWAT) and play an essential role in browning during cold exposure^58^. As such, we sought to investigate if mice receiving Sb-Exe4 displayed enhanced UCP1-mediated thermogenesis. While we did not observe a significant difference in adipose weight, we observed a tendency in reduced adipose tissue mass from Sb-Exe4 mice over Sb-Empty mice (Supplementary Figure S7 and Supplementary Figure S8). The browning of sWAT, as evidenced by emerge of beige adipocytes within sWAT, was observed under cold conditions (Supplementary Figure S7F). In addition, the adipocyte cell size of sWAT from Sb-Exe4 mice was significantly reduced compared to that of Sb-Empty mice (Supplementary Figure S7G).

## Discussion

*S. boulardii* has caught a lot of attention as an AMT chassis for its eukaryotic characteristics^33,34^. However, there is limited evidence of *S. boulardii* as an AMT chassis, particularly in the field of metabolic disorders. In this study, we investigated the biological efficiency of a genetically modified *S. boulardii* secreting well-established GLP-1R agonists. First, we demonstrated that *S. boulardii* can be engineered to secrete high levels of GLP-1 and Exendin-4 aerobically and anaerobically. While both Sb-GLP1 and Sb-Exe4 produced similar titres of the GLP1-R agonist, only Sb-Exe4 successfully stimulated the insulin secretion from primary pancreatic islets of wild-type mice (Figure 1). The substantial difference between the two GLP-1R agonists could be explained by the significant difference in peptide stability^17^, hence more prolonged exposure to the GLP1-R agonist and a higher likelihood of activating insulin secretion. The fold increase of insulin secretion is in a similar range as previously been reported with microbial-produced GLP-1R agonists^27^. No significant effect was observed in islets of GLP-1R knockout mice, indicating that the insulin-stimulating response from the Sb-Exe4 was dependent on the GLP-1R.

We next sought to explore the biological efficiency of *in situ* biosyntheses of Exendin-4 by *S. boulardii* in a mouse gastrointestinal tract. Although *in situ* biosyntheses of GLP-1 in animals have previously been reported with other bacterial probiotics^27–29,31^, we demonstrated *in situ* production of Exendin-4 with a eukaryotic host and identified a synergistic effect of cold exposure and Sb-Exe4. We did not observe any significant effect of Sb-Exe4 at room temperature, which could be explained by the *S. boulardii*’s inherent anti-obesity properties against obese mice^32^ or the faster transient time in mice^47^ resulting in insufficient exposure of Exendin-4; however, we observed a drastic effect during cold exposure. Exposing mice to cold has been observed to affect the circulating levels of various hormones affecting whole-body metabolism and energy expenditure, such changes trigger an increase in food intake^59,60^. As such, *in situ* delivery of Exendin-4 in the gastrointestinal tract may interact with the GLP-1R, hence suppressing the increased food intake induced by cold. Therefore, oral administration of Sb-Exe4 and cold exposure resulted in a more robust suppression of food intake and body weight than at room temperature. Moreover, the Sb-Exe4 mice exhibited an increased activity and decreased energy expenditure during cold exposure, suggesting that the mice might acquire a compensatory ability to preserve their energy in the cold environment.

Receiving the Sb-Exe-4 consistently decreased the RER compared to the Sb-Empty treated mice. These results are consistent with previous reports on Exendin-4^61^ and other GLP-1R agonists^62^. The lower RER is also in line with the increased expression of fatty acid transporter *Cd36* in the liver, showing higher mobilisation of fatty acids in the animal body. The activation of *Cd36* has also previously been demonstrated to protect mice from lower blood sugar^63^. In addition, we observed an increased blood circulation of glucagon and hepatic expression of the gluconeogenic genes *G6pc* and *Pepck* in cold exposed and Sb-Exe4 treated mice, suggesting that the decreased food intake might also result in a more active gluconeogenesis^53^. The induced hepatic expression of *Pepck* has previously been reported with a microbial-produced GLP-1R agonist in high-fat fed mice^27^. Moreover, the combined effect of cold exposure and Sb-Exe4 also showed a consistent tendency of reduced white adipose tissue mass and reduced adipocyte cell size of sWAT. The change in the adipose tissue morphology is in line with the reduced circulating TNF-α levels that have been shown to correlate with the degree of adiposity and chronic inflammation^64,65^. These observations on adipose development are consistent with previous reports on Exedin-4 and other GLP-1R agonists^61,62^.

Exploring novel delivery strategies for effective therapeutic compounds and peptides is of great interest for improving patient adherence and treatment. Our study has demonstrated that *S. boulardii* is an attractive AMT chassis able to deliver therapeutic peptides to potentially improve the treatment of metabolic disorders. Several challenges remain for oral peptide delivery by AMTs before being applied in a clinical setting. However, here we present the first step towards a new platform for oral delivery of therapeutic peptides, which holds the promise to significantly improve the treatment quality of patients.

## Methods

### Strains and plasmid construction

Plasmids, strains, primers, and sequences used in this study are listed in Supplementary Table S1, S2, S3 and S4. Oligonucleotides and gBlocks were ordered from Integrated DNA Technologies (IDT). All integration plasmids used in this study were linearised with the restriction enzyme SfaAI (FastDgiest Enzyme, Thermo Scientific™) and BcuI (FastDgiest Enzyme, Thermo Scientific™) gel purified (Thermo Scientific™ GeneJET Gel Purification Kit). All plasmid assemblies were conducted with Gibson Assembly^66^ and transformed into One Shot® TOP10 *Escherichia coli* (Thermo Fisher Scientific). All *E. coli* were grown in lysogeny broth (LB) media containing 5 g/L yeast extract, 10 g/L tryptone and 10 g/L NaCl; (Sigma Aldrich) supplemented with 100 mg/L ampicillin sodium salt (Sigma Aldrich). LB agar plates contained 1% agar (Sigma Aldrich). *S. boulardii* with uracil auxotroph was obtained from our previous work^47^. The uracil auxotroph was used for all experiments. *S. boulardii* transformations were performed via high-efficiency yeast transformation using the LiAc/SS carrier DNA/PEG method^67^. Genomic integrations cassettes were digested with the restriction enzyme NotI (FastDgiest Enzyme, Thermo Scientific™) prior to transformation and transformed with various helper plasmids and pre-expressed Cas9 from pCfB2312^68^. All transformations were heat-shocked in a water bath at 42°C for 60 min. A recovery step was included in the transformation protocol. The transformation tubes were microcentrifuge for 2 min at 3000 g. Pellets were resuspended in 500 µL of YPD and incubated for 2 – 3 hours. All yeast transformations were plated on synthetic complete (SC) plates containing 1.7 g/L yeast nitrogen base without amino acids and ammonium sulphate (Sigma Aldrich, Merck Life Science), 1 g/L monosodium glutamate (Sigma Aldrich, Merck Life Science), 1.92 g/L Yeast Synthetic Drop-out Medium Supplements without uracil (Sigma Aldrich) and 200 mg/L geneticin (G418; Sigma Aldrich) at 37°C. Colony-PCR using OneTaq (Thermo Scientific™) was used to confirm the genomic integration. Primers flanking the integration were used to confirm the integration. Genomic DNA was extracted by boiling cells at 70°C for 30 min in 20 mM NaOH. One single amplification band, ∼2000 bp, on gel electrophoresis indicated a successful integration into both chromosome copies. The strains were cured for pCfB2312 and helper plasmids after genome integration.

EasyClone MarkerFree vectors pCfB2899, pCfB2904, pCfB2909, pCfB3035 and helper plasmids pCfB6910, pCfB6912, pCfB6915, pCfB6920 and pCfB10292 were adapted as previously described^68^. The nourseothricin selection marker (NatMX) was replaced with a uracil selection marker (*URA3*).

All *S. boulardii* were cultured in yeast extract peptone dextrose (YPD) media containing 10 g/L yeast extract, 20 g/L casein peptone and 20 g/L glucose (Sigma Aldrich) at 37°C. Strain selection was done on YPD plates containing 20 g/L agar or minimal synthetic media (DELFT) containing 7.5 g/L (NH_4_)_2_SO_4_, 14.4 g/L KH_2_PO_4_, 0.5 g/L MgSO_4_ x 7H_2_O, 20 g/L glucose, 2 mL/L trace metals solution, and 1 mL/L vitamins^69^. The pH was adjusted to 6.

### ELISA

*S. boulardii* cultures were incubated in 2 mL DELFT medium supplemented with 20 mg/L uracil in a 24-deep well plate (Axygen®, VWR) with a sandwich cover (Enzyscreen). Aerobic cultivation with an initial OD_600_ of 0.05 was performed with continuous shaking at 250 RPM at 37°C. Anaerobic cultivation with an initial OD_600_ of 0.05 was performed without shaking at 37°C in media reduced in the anaerobic coy chamber (gas mixture, 95% N_2_ and 5% H_2_) for two days prior to use. All cultures were harvested after 24 hours. Cell cultures were spun down at 10,000 g for 10 min at 4°C. GLP-1 secreted was quantified with Human GLP-1 ELISA (Abcam Cat. #ab184857), and Exendin-4 secreted was quantified with Exendin-4 EIA (Phoenix Cat. #EK-070-94). The signals were detected by OD_450_ using a microplate reader Synergy™ H1 BioTek.

### Glucose-stimulated insulin release from isolated islets (GSIS)

Islets were isolated from C57BL/6 wild-type and C57BL/6 GLP-1R knockout mice using collagenase perfusion as previously described^70^. After the isolation, islets were incubated in RPMI-1640 medium overnight. Islets were washed in Krebs Ringer Hepes buffer (KRH) with 2.8 mM glucose. Five islets per group were incubated with 16.7 mM glucose KRH, with either 2 μM Exendin-4 (positive control) or cell-free supernatants from Sb-Empty, Sb-GLP-1 or Sb-Exe4 cultures diluted 1:3 at 37°C for 2 hours. After incubation, islets were sedimented, and insulin was measured in the supernatants using the insulin ELISA kit (Crystal Chem). All stimulations were performed in duplicates.

### Animal studies

All animal experiments were conducted according to the Danish guidelines for experimental animal welfare, and the study protocols were approved by the Danish Animal Experiment Inspectorate (license number 2020-15-0201-00405). The study was carried out in accordance with the ARRIVE guidelines^71^. All *in vivo* experiments were conducted on male C57BL/6NTac mice (6-8 weeks old; Taconic Bioscience). Unless otherwise stated, all mice were housed at room temperature on a 12-hour light/dark cycle and given *ad libitum* access to water and a high-fat diet (Research Diet, D12451 Rodent Diet with 45 kcal% fat). The mice received the high-fat diet from day 0 and were renewed once a week. The mice were orally administered via intragastric gavage with ∼10^8^ CFU of *S. boulardii* (either Sb-Empty or Sb-Exe4) in 100 µL of 1x PBS and 20% glycerol at 01:00 p.m. unless otherwise stated. The mice were randomised according to body weight and acclimated for at least one week prior to oral administration. The researchers were blinded in all mouse experiments.

The mice were euthanised after a two to four hours fast, by anaesthesia (25% hypnorm, 25% dormicum in sterile water, (0.01 mL/g mouse)) followed by blood collection from the vena porta and heart, and cervical dislocation. Plasma was separated in blood collection tubes containing a PST™ plasma separator gel, lithium heparin additive (BD Microtainer®) and 30 µL of 5x cOmplete™ Protease Inhibitor Cocktail (Roche Applied Science Cat. #04693116001) and 0.5 mg/mL Sitagliptin Phosphate (Sigma Aldrich Cat. #PHR1857). The weight of subcutaneous and epididymal weight adipose tissue and interscapular brown adipose tissue was measured and snap-frozen in liquid nitrogen for further processing. The left lobe of the liver was dissected and snap-frozen in liquid nitrogen for further processing. All samples were stored at -80°C.

### Synergetic effect of Sb-Exe4 and cold exposure on energy homeostasis

The mice were divided into two groups and were orally administered for 29 days with either Sb-Empty (n = 9) or Sb-Exe4 (n = 9). Body weight was monitored weekly. Faeces was collected on day 0, 7, 14, and 21. The mice were moved into metabolic cages from day 29. On day 24 the mice were exposed to cold (8°C). The mice were euthanised on day 29.

### Metabolic cages

Indirect calorimetry was performed using the Home Cage System (TSE systems). The mice were acclimated to the TSE cabinets for three days before measurements. Gas exchanges and food intake were recorded every 20 minutes. The cages were put in climate-controlled rodent incubators (PhenoMaster Climate Chamber, TSE systems) set to 22°C for room temperature monitoring and 5°C (measured 8°C) for the cold exposure monitoring. All the metabolic phenotyping data were analysed by averaging data from three to eight days of measurements.

### CFU assessment in the faecal and gut samples

The faeces and gut matter were collected in pre-weighed 1.5 mL or 2.0 mL Eppendorf tubes containing 1 mL of 1x PBS and 50% glycerol and weighed again to determine the faecal weight. All sample preparation for assessing CFU numbers was kept at 4°C and followed the same practice. The faecal samples were homogenised by vortexed at ∼2400 rpm for 20 min. The samples were then spun down at 100 g for 30 seconds, followed by a dilution series, where 5 µL of each dilution was plated in duplicates or triplicates. The faecal samples were plated on SC supplemented with 20 mg/L uracil plates containing 100 mg/L ampicillin, 50 mg/L kanamycin, and 30 mg/L chloramphenicol (Sigma Aldrich, Merck Life Science) for preventing bacterial growth and 1 g/L 5-Fluoroorotic Acid (5-FOA; Nordic BioSite) for preventing other yeast species from growing.

### Cytokine and hormone analysis

Plasma samples from mice were analysed with Mesoscale U-PLEX^®^ multiplex custom assays according to the manufacturer’s instructions.

### Real-Time Quantitative PCR

The tissues were homogenised in TRI reagent (Sigma Aldrich), and RNA was isolated with Direct-zol™ RNA Miniprep Plus (Zymo Research), including a DNase I treatment. The RNA concentration and quality were determents using a NanoDrop 1000 spectrophotometer. Reverse transcriptomics to cDNA was performed with High-Capacity cDNA Reverse Transcription Kit (Thermo Scientific™). mRNA expression levels were determined by qPCR. 2x SensiFAST SYBR® Lo-ROX mix (Nordic BioSite) was used for all qPCR reactions, consisting of 5 µL of master mix, 0.03 µL of each primer (100 µM, see Supplementary Table 3) and 5 µL of 20 to 25 ng of cDNA. Low quality RNA samples were removed for further analysis. mRNA expression levels in sWAT and iBAT samples were normalised to the mean of *Tbp* and *Tfiib* expression. Liver samples were normalised to *Hprt* and *Tfiib* expression in the conventional mice. All qPCR reactions followed the same thermocycler program that consisted of an initial 3 min step at 98°C, followed by 40 cycles of 98°C for 10 s, 60°C for 15 s, and 72°C for 30 s, along with a final melting curve consisting of a single cycle of 98°C for 5 s, 60°C for 1 min, 98°C for 5 s. All samples were run in technical duplicate. All qPCR runs were performed on the Roche LightCycler® 480 Real-time PCR System in LightCycle® 480 Multiwell Plate 384, clear plates (Roche).

### Statistical testing

All statistical analysing were performed in RStudio version 4.1.0 with the rstatix and DescTools package. Unless otherwise noted, all data are presented as means ± SEM. The statistical significance level was set at p < 0.05. The normality of the data was checked using the Shapiro-Wilk test. Statistical differences between groups of two were analysed with a dependent sample t-test or Wilcoxon singed-rank test. Bonferroni adjustments were used for multiple comparison (e.g., groups and genes). Comparison of three or more groups were analysed by ANOVA with a Dunnett’s post hoc test.

## Data availability

All data generated or analysed during this study are included in this published article and its Supplementary Information files. Further inquiries can be directed to the corresponding author/s.

## Acknowledgement

This work received funding from The Novo Nordisk Foundation under NNF grant number: NNF20CC0035580, NNF Challenge programme CAMiT under Grant agreement: NNF17CO0028232 and the European Union’s Horizon 2020 research and innovation programme under the Marie Skłodowska-Curie Grant agreement No 813781. Fredrik Bäckhed is the Torsten Söderberg Professor in Medicine and a Wallenberg Scholar. We are grateful to David Romero Suarez and Vakil Takhaveev for the initial discussion on strain engineering. Thank you Philip Tinggaard Thomesen for providing the pCfB6910, pCfB6912, pCfB6915, pCfB6920 and pCfB10292 gRNA plasmids. We are also grateful to Susanne Kammler and DTU Bio Facility’s animal caretakers, Kenneth Rene Worm, Maja Danielsen and Heidi Arps, for assistance in carrying out some of the *in vivo* experiments.

## AUTHOR INFORMATION

### Contributions

All authors contributed in conceived the study and contributed to the design of the study. K.A.H. carried out all the strain engineering and characterisation. T.U.G. carried out the GSIS assay. K.A.H., H.Z., V.K., V.E.R., and R.V.U. carried out all the *in vivo* characterisations. K.A.H. and V.K conducted the qPCR assay. H.Z. carried out the histology assay. V.R. carried out the western blot. T.U.G. carried out the mesoscale-discovery assay. K.A.H. analysed the data and wrote the manuscript. F.B., T.U.G., R.V.U, and M.O.A.S. supervised the study. All authors contributed to the discussion of the results. All authors read and approved the final manuscript.

### Corresponding authors

Correspondence to Morten Otto Alexander Sommer and Ruben Vazquez-Uribe

