## Supplementary Information for "Oral delivery of GLP-1R agonist by an engineered probiotic yeast strain has anti-obesity effects in mice"

#### Supplementary Figures:

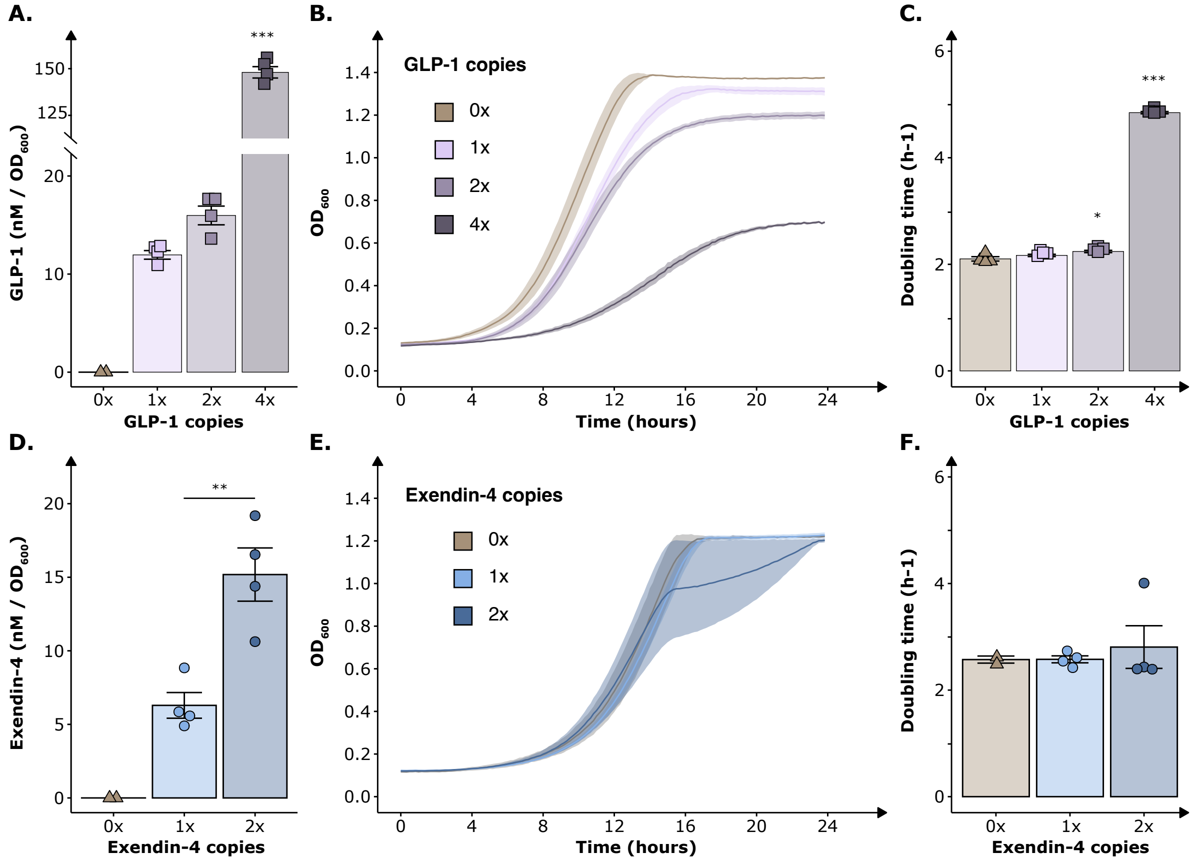


**Figure S1. *In vitro* characterisation of the *S. boulardii* producer strains. (A)** Concentration of GLP-1 normalised to $OD_{600}$ in the supernatant from *S. boulardii* with different copy number of gene cassettes expressing GLP-1. (**B)** Real-time $OD_{600}$ measurement of the different GLP-1 producer strains. (**C)** Doubling time for the different GLP-1 producer strains. (**D)** Concentration of Exendin-4 normalised to $OD_{600}$ in the supernatant from *S. boulardii* with different copy number of gene cassettes expressing Exendin-4. (**E)** Real-time $OD_{600}$ measurement of the different Exendin-4 producer strains. **(F)** Doubling time for the different Exendin-4 producer strains. Data presented as mean ± SEM (n = 2 - 4). * $p < 0.05$, ** $p < 0.01$, *** $p < 0.001$. A, B and F were analysed with one-way ANOVA, Dunnett's post hoc test, with 1x as reference in A and 0x as reference in C and F. D was analysed by dependent sample t-test.

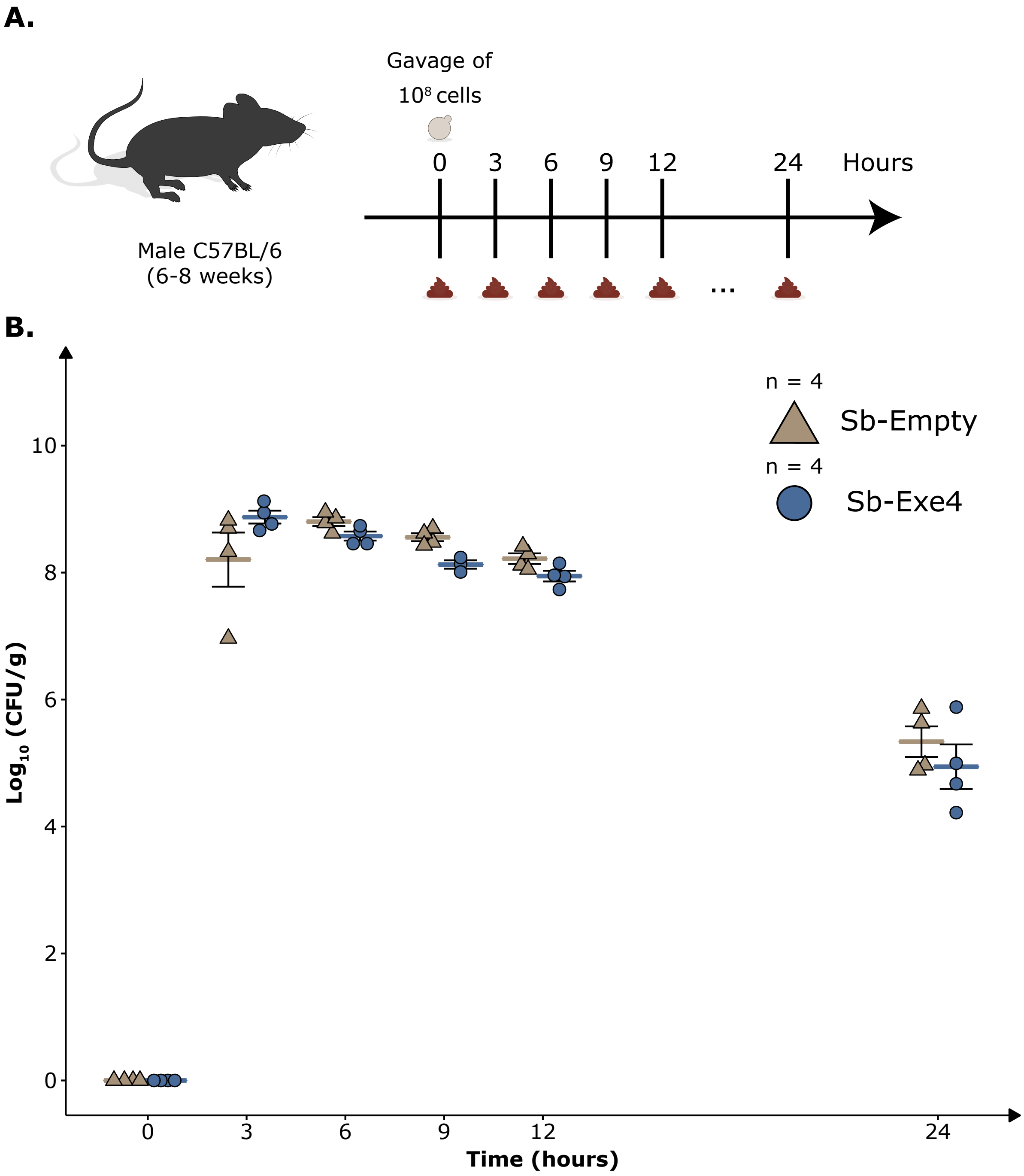


**Figure S2. Characterisation of colonisation in conventional mice over 24 hours. (A)** Schematic overview of the study plan. Male C57BL/6 mice were divided into two groups, orally administered with either Sb-Empty or Sb-Exe4 for 5 days to ensure steady engraftment. (**B**) Abundance (CFU/g faeces) of Sb-Empty and Sb-Exe4 at 3, 6, 9, 12, and 24 hours after the last oral administration of Sb-Empty and Sb-Exe4 in the mice. Data presented as mean ± SEM (n = 4). Each point represents CFU/g faeces in one mouse. Differences were analysed with two-way ANOVA using the factors time and strain.

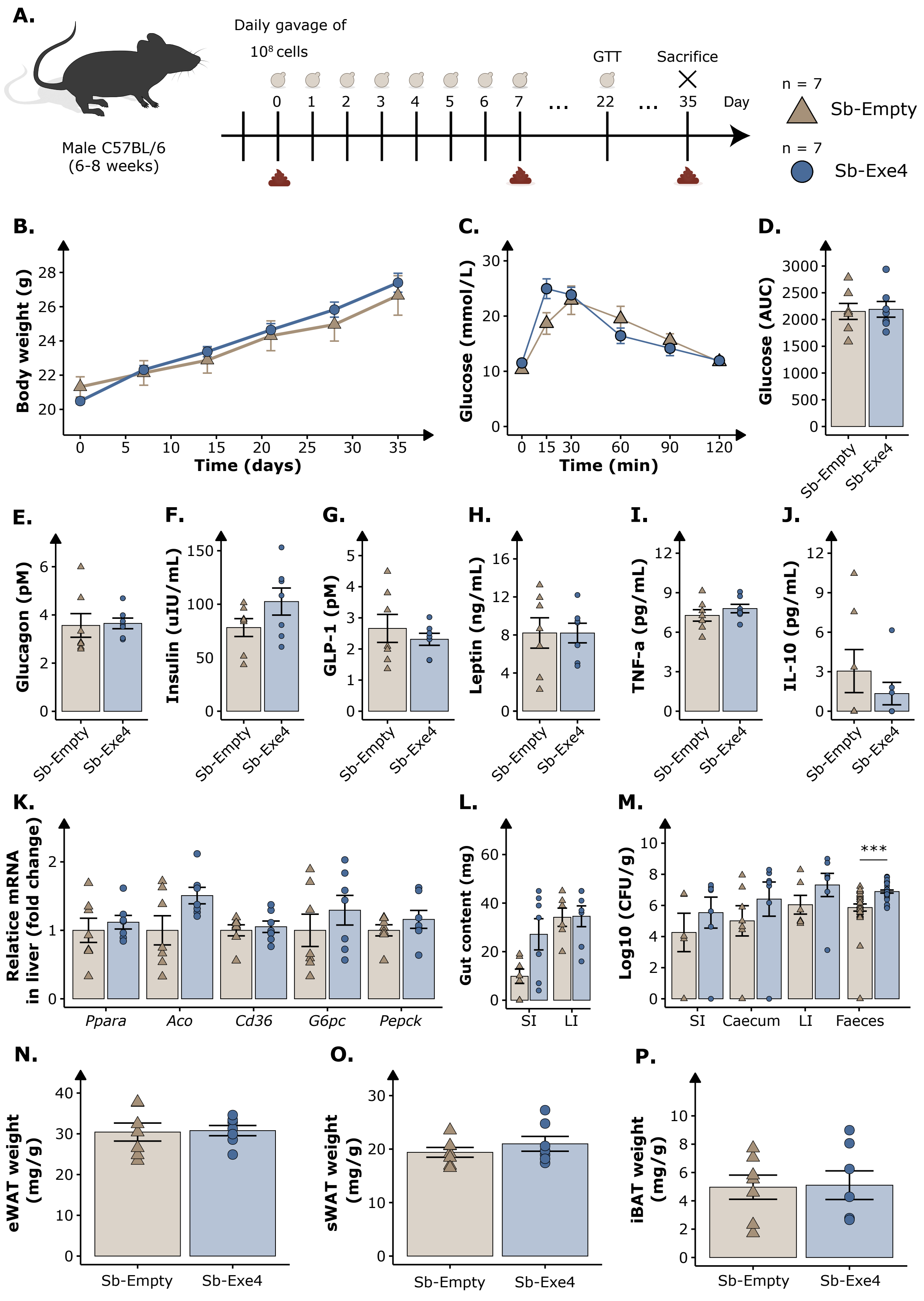


**Figure S3.** ***In vivo*** **characterisation of Sb-Exe4 in male C57BL/6 mice for 35 days. (A)** Schematic overview of the study plan. Male C57BL/6 mice were divided into two groups, orally administered with either Sb-Empty or Sb-Exe4. Mice were initiated on a 45 % kcal high-fat diet on day 0. **(B)** Body weight was monitored weekly for 35 days while the mice were fed high-fat diet and oral administered their respective strain (n = 7). **(C)** Glucose challenge was performed with an intraperitoneal glucose tolerance test (IP-GTT) 22 days after the first oral administration (n = 7). **(D)** Mean total area under the curve (AUC) of the IP-GTT (n = 7). Fasting vena cava levels of **(E)** glucagon (pM), **(F)** insulin (uIU/mL), **(G)** GLP-1 (pM), **(K)** leptin (ng/mL), **(L)** TNF-α (pg/mL), and **(M)** IL-10 (pg/mL) at the end of the study quantified with Meso Scale Discovery (n = 7). **(K)** Relative mRNA of *Ppara*, *Aco*, *Cd36*, *G6pc* and *Pepck* in the liver of the mice at the end of the study. Data presented as fold change normalised to Sb-Empty (n = 5 – 7). **(L)** Weight (mg) of gut content collected from small intestine (SI) and large intestine (LI) from the mice at the end of the study (n = 6 – 7). **(M)** Abundance ($Log_{10}$ CFU/g faeces) of Sb-Empty and Sb-Exe4 in SI (n = 5 – 7), caecum (n = 7), LI (n = 6 – 7), and faeces of the mice (n = 32 – 33). **(N)** Epididymal white adipose tissue (eWAT; n = 7), **(O)** subcutaneous white adipose tissue (sWAT; n = 7), and **(P)** interscapular brown adipose tissue (iBAT; n = 7) weight (mg) normalised to body weight (g) of each mouse at the end of the study. Data presented as mean ± SEM. P-value written out $p < 0.1$, * $p < 0.05$, ** $p < 0.01$, $p < 0.001$. B and C were analysed with two-way ANOVA using the factors time and strain. D – Q were analysed with Wilcoxon signed-rank test with Bonferroni adjustment for multiple comparison.

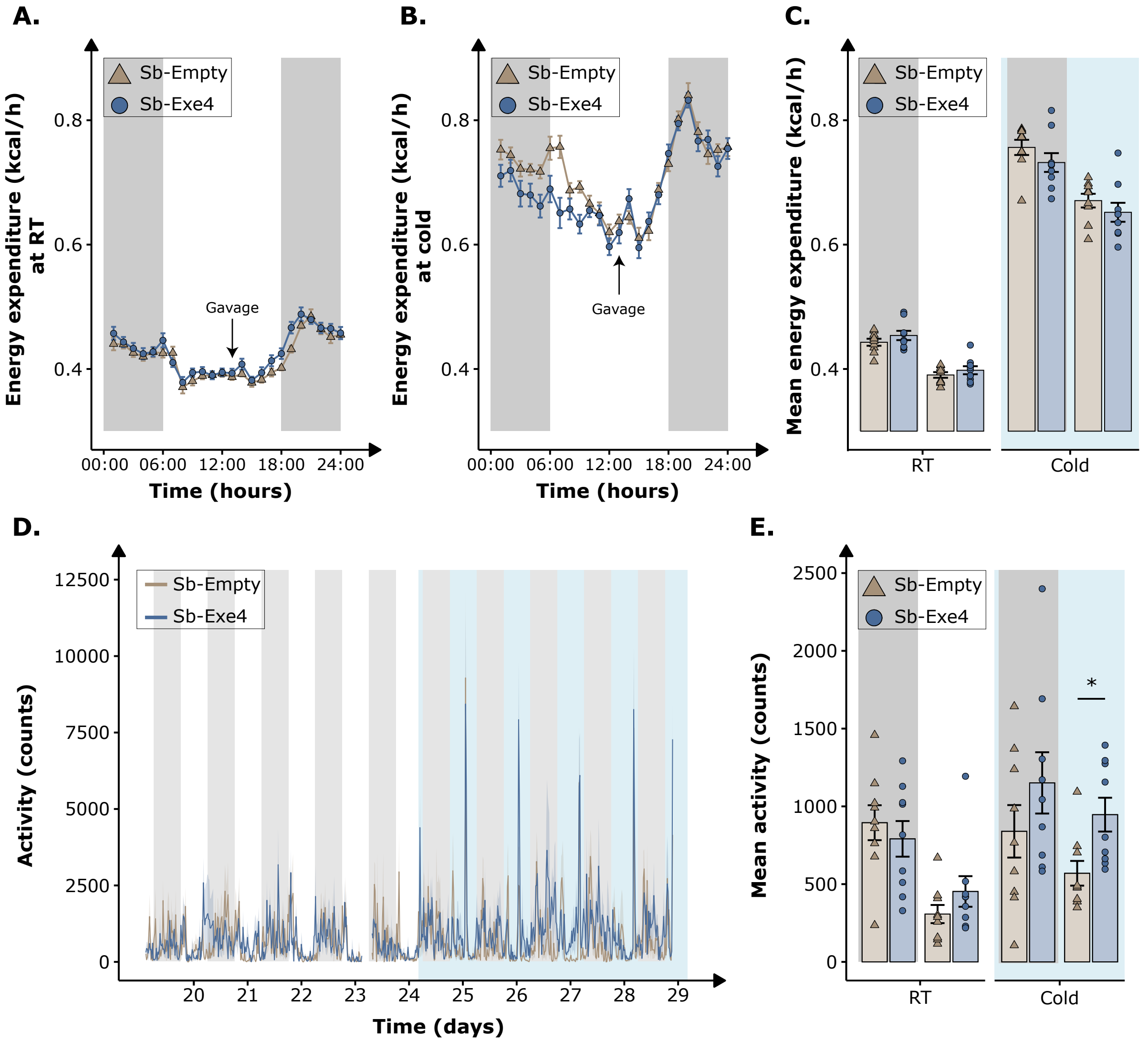


**Figure S4. Synergetic effect of Sb-Exe4 and cold exposure on energy homeostasis in male C57BL/6 mice fed a high-fat diet for 29 days. (A)** Mean hourly energy expenditure (kcal/h) under a 12:12 light-dark cycle over a 5-day period at room temperature (RT; 22 °C) and **(B)** a 5-day period at cold (8 °C). **(C)** Mean daily energy expenditure during light and dark cycle at RT and cold. **(D)** Activity (counts) under a 12:12 light-dark cycle over an 8-day period. **(E)** Mean daily activity during light and dark cycle at RT and cold. Black shaded area indicates dark period. Light blue shaded area indicates cold exposure (8 °C). Data presented as mean ± SEM (n = 9). * $p < 0.05$. All samples were analysed by Wilcoxon signed-rank test with Bonferroni adjustment for multiple comparison.

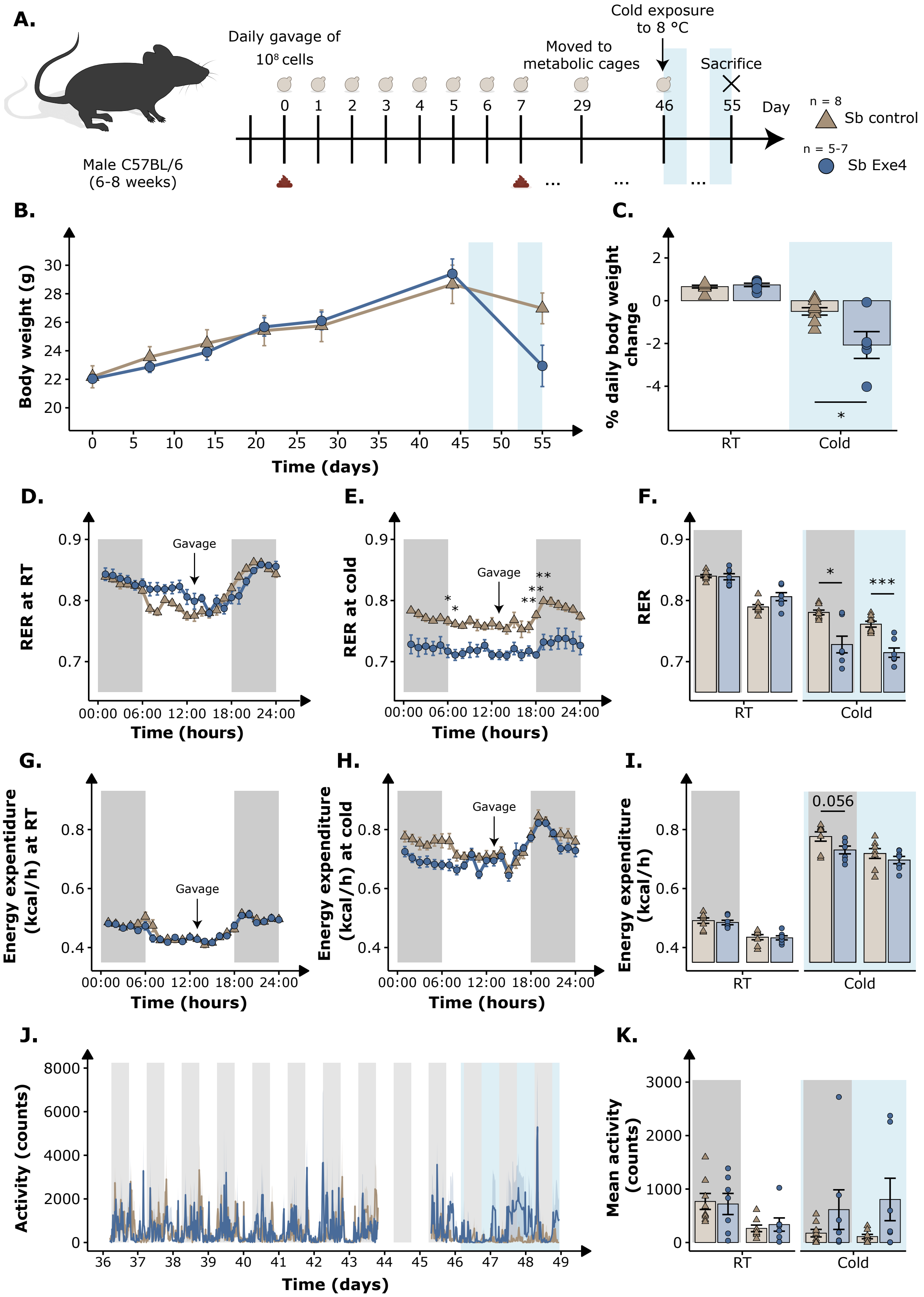


**Figure S5. Synergetic effect of Sb-Exe4 and cold exposure on energy homeostasis in male C57BL/6 mice fed a high-fat diet for 55 days. (A)** Schematic overview of the study plan. Male C57BL/6 mice were divided into two groups, orally administered with either Sb-Empty or Sb-Exe4. Mice were initiated on a 45 %kcal high-fat diet on day 0. **(B)** Body weight was monitored weekly for 55 days while the mice were fed high-fat diet and orally administered their respective strain (n = 5 – 8). **(C)** Mean daily body weight change at room temperature (RT; 22 °C) and cold (8 °C). **(D)** Mean hourly respiratory exchange ratio (RER; $VCO_2$ / $VO_2$) under a 12:12 light-dark cycle over an 8-day period at RT and **(E)** a 3-day period at cold. **(F)** Mean RER during light and dark cycle at RT and cold. **(G)** Mean hourly energy expenditure (kcal/h) under a 12:12 light-dark cycle over an 8-day period at RT and **(H)** a 3-day period at cold. **(I)** Mean energy expenditure during light and dark cycle at RT and cold. **(J)** Activity (counts) under a 12:12 light-dark cycle over an 11-day period. **(K)** Mean daily activity during light and dark cycle at RT and cold. Data are presented as the mean ± SEM (n = 7 – 8). Black shaded area indicates dark period. Light blue shaded area indicates cold exposure (8 °C). P-value written out $p < 0.1$, * $p < 0.05$, ** $p < 0.01$. All samples were analysed by Wilcoxon signed-rank test with Bonferroni adjustment for multiple comparison.

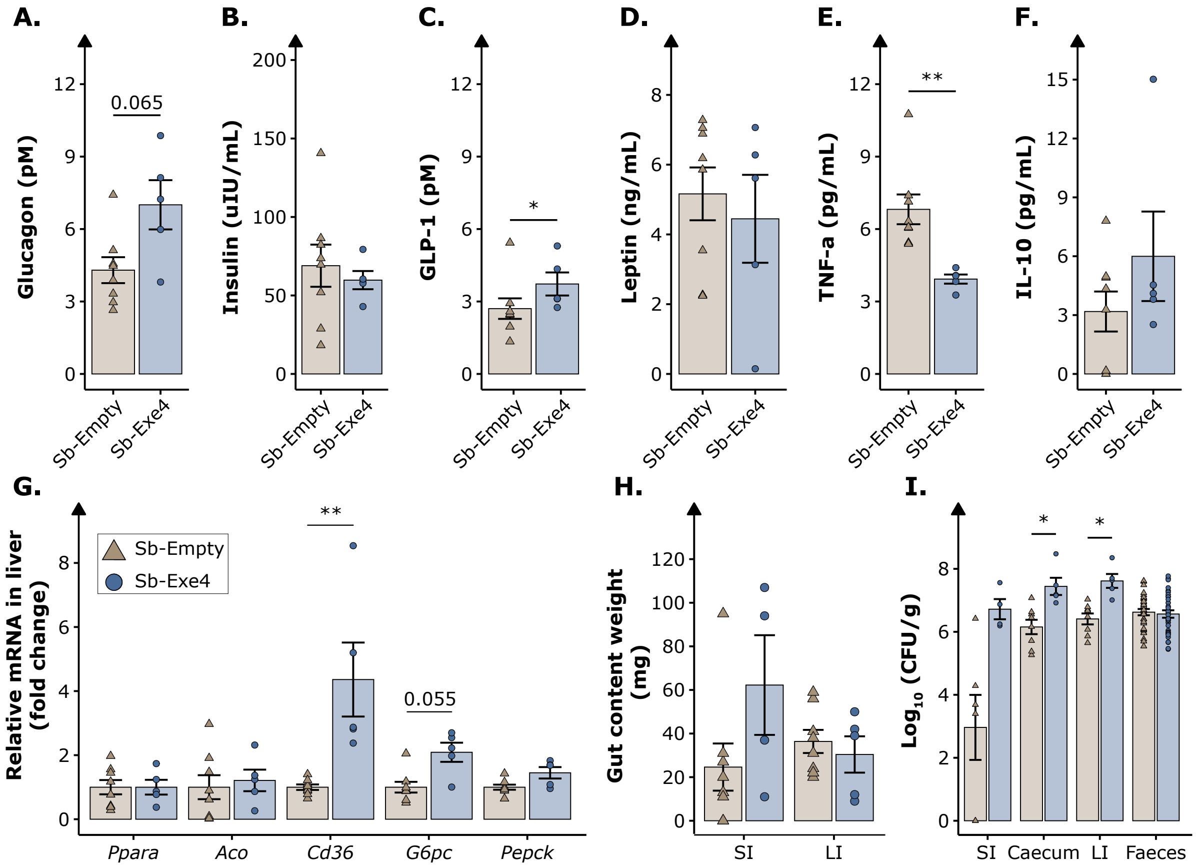


**Figure S6. End-point characterisation of the synergistic effect of cold exposure and Sb-Exe4 in male C57BL/6 mice for 55 days.** Fasting vena cava levels of **(A)** glucagon (pM), **(B)** insulin (uIU/mL), **(C)** GLP-1 (pM), **(D)** leptin (ng/mL), **(E)** TNF-α (pg/mL) and **(F)** IL-10 (pg/mL) at the end of the study quantified with Meso Scale Discovery (n = 5 – 8). **(G)** Relative mRNA of *Ppara*, *Aco*, *Cd36*, *G6pc* and *Pepck* in the liver at the end of the study. Data presented as fold change normalised to Sb-Empty (n = 5 – 8). **(H)** Weight (mg) of gut content collected from small intestine (SI) and large intestine (LI) of the mice (n = 4 – 8). **(I)** Abundance ($Log_{10}$ CFU/g faeces) of Sb-Empty and Sb-Exe4 in SI (n = 4 – 6), caecum (n = 5 – 8), LI (n = 5 – 8), and faeces (n = 31 – 32) in the mice. Data are presented as the mean ± SEM. P-value written out < 0.1, * p < 0.05, ** p < 0.01. All data were analysed by Wilcoxon signed-rank. Bonferroni adjustments were used for multiple comparison.

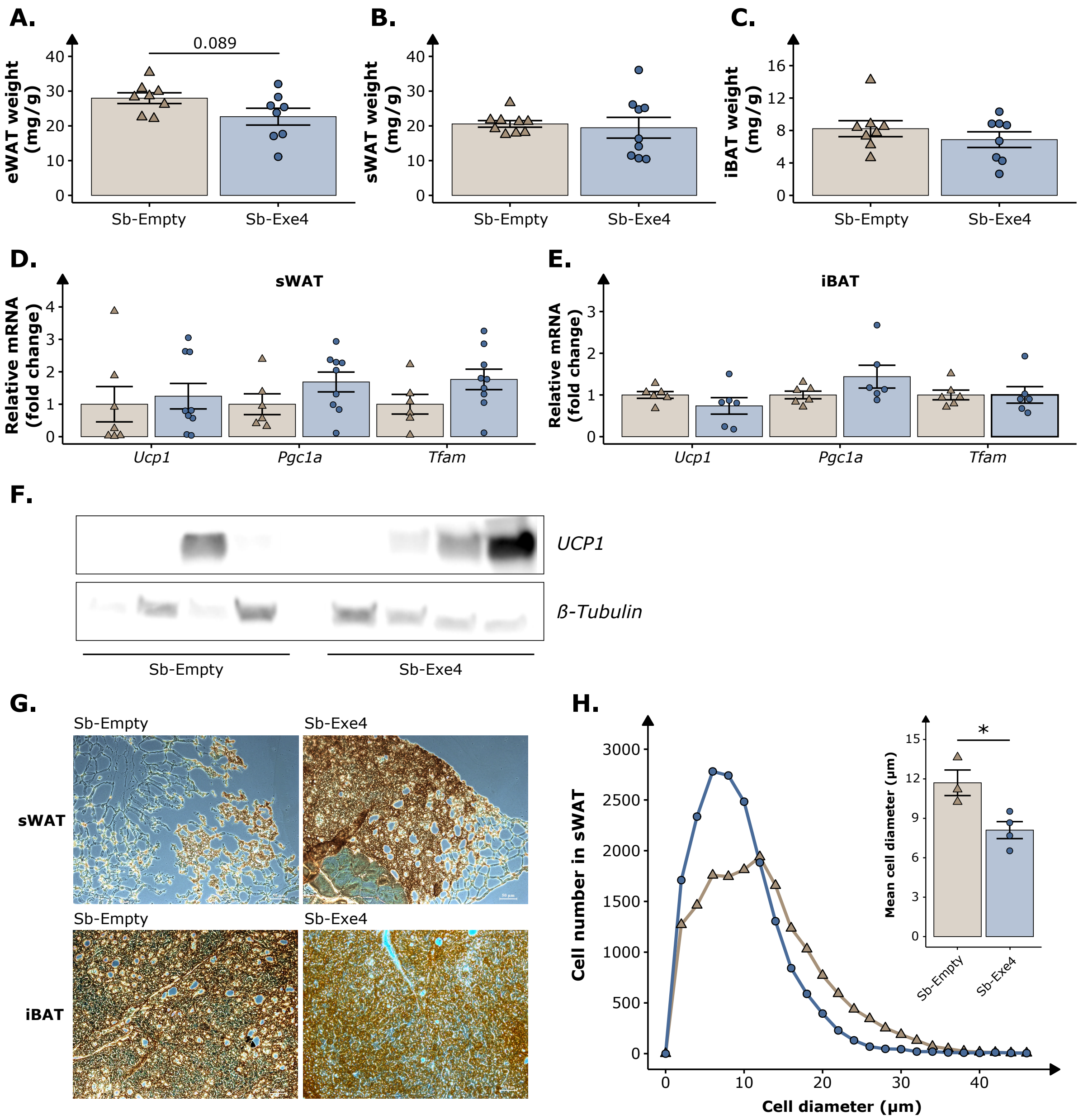


**Figure S7. End-point characterisation of the synergistic effect of cold exposure and Sb-Exe4 on the adipose tissue in male C57BL/6 mice for 29 days. (A)** Epididymal white adipose tissue (eWAT; n = 7 – 8), **(B)** subcutaneous white adipose tissue (sWAT; n = 9), and **(C)** interscapular brown adipose tissue (iBAT; n = 8) weight (mg) normalised to body weight (g) of each mouse at the end of the study. **(D)** Relative mRNA of *Ucp1*, *Pgc1a,* and *Tfam* in sWAT at the end of the study. Data presented as fold change normalised to Sb-Empty (n = 6 – 9). **(E)** Relative mRNA of *Ucp1*, *Pgc1a,* and *Tfam* in iBAT at the end of the study. Data presented as fold change normalised to Sb-Empty (n = 7 – 8). **(F)** Western blot of UCP1 expression in sWAT (n = 4). **(G)** Immunostaining of UCP1 in sWAT and iBAT sections. The brown colour represents the presence of UCP1 protein. Bar: 50 µm. **(H)** The distribution of adipocyte cell diameter and cell number on the sWAT sections quantified by ImageJ (n = 3 – 4). Data are presented as the mean ± SEM. * $p < 0.05$. A-C, G were analysed by dependent sample t-test. D-E were analysed by Wilcoxon signed-rank test. Bonferroni adjustments were used for multiple comparison.

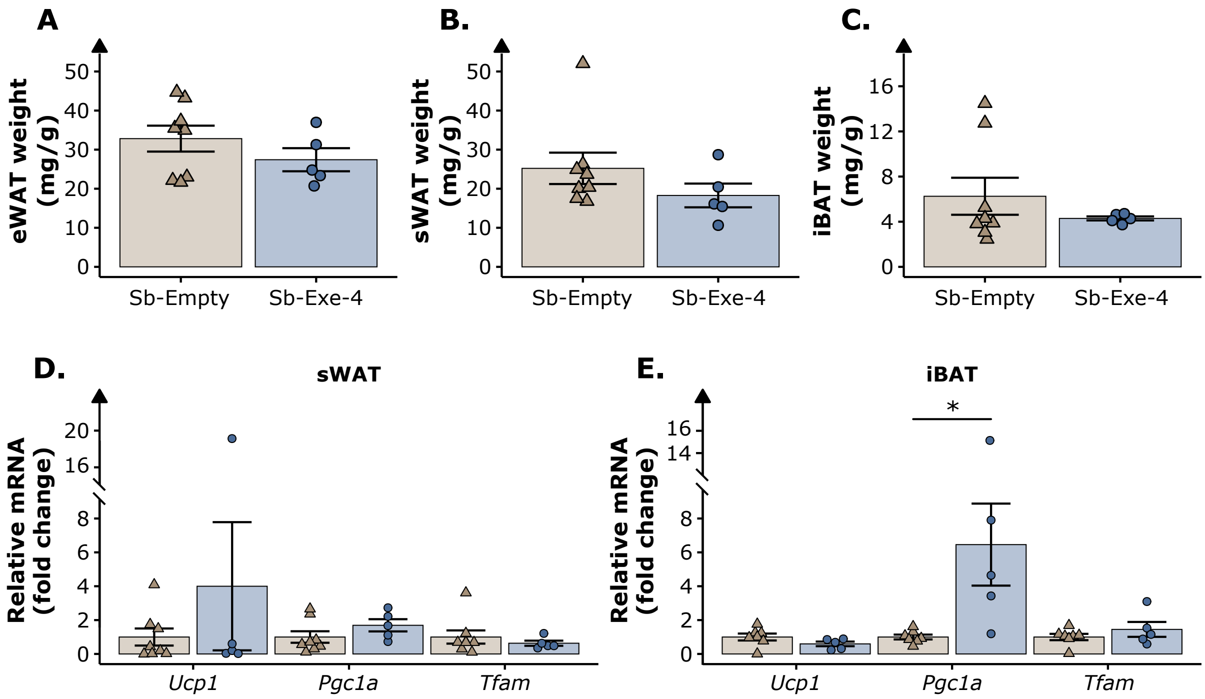


**Figure S8. End-point characterisation of the synergistic effect of cold exposure and Sb-Exe4 on the adipose tissue in male C57BL/6 mice for 55 days. (A)** Epididymal white adipose tissue (eWAT), **(B)** subcutaneous white adipose tissue (sWAT), and **(C)** interscapular brown adipose tissue (iBAT) weight (mg) normalised to body weight (g) of each mouse at the end of the study. **(D)** Relative mRNA of *Ucp1*, *Pgc1a,* and *Tfam* in sWAT at the end of the study. Data presented as fold change normalised to Sb-Empty. **(E)** Relative mRNA of *Ucp1*, *Pgc1a,* and *Tfam* in iBAT at the end of the study. Data presented as fold change normalised to Sb-Empty. Data are presented as the mean ± SEM (n = 5 – 8). * $p < 0.05$. A-C were analysed by dependent sample t-test. D-E were analysed by Wilcoxon signed-rank test. Bonferroni adjustments were used for multiple comparison.

### Supplementary Tables

**Table S1. Plasmids used in this study**

| Plasmid name | Genotype | Marker (*E. coli / S. boulardii)* | Reference |
|---|---|---|---|
| pCfB2312 | CEN6_ARS4 Amp $P_{TEF1}$-Cas9-$t_{CYC1}$ kanMX | Amp / KanMX | [1] |
| pCfB2899 | X-2-MarkerFree | Amp | [1] |
| pCfB2899-GLP1 | pCfB2899: $P_{TDH3}$-GLP1-$t_{DIT1**}$ | Amp | This study |
| pCfB2904 | XI-3-MarkerFree | Amp | [1] |
| pCfB2904-GLP1 | pCfB2904: $P_{TDH3}$-GLP1-$t_{DIT1**}$ | Amp | This study |
| pCfB2904-Exe4 | pCfB2904: $P_{TDH3}$-Exe4-$t_{DIT1**}$ | Amp | This study |
| pCfB2909 | XII-5-MarkerFree | Amp | [1] |
| pCfB2909-GLP1 | pCfB2909: $P_{TDH3}$-GLP1-$t_{DIT1**}$ | Amp | This study |
| pCfB2909-Exe4 | pCfB2909: $P_{TDH3}$-Exe4-$t_{DIT1**}$ | Amp | This study |
| pCfB3035 | X-4-MarkerFree | Amp | [1] |
| pCfB3035 | pCfB3035: $P_{TDH3}$-GLP1-$t_{DIT1**}$ | Amp | This study |
| pCfB6910 | pCfB3020-URA; $P_{SNR52}$-gRNA(X-2)-$t_{SUP4}$ | Amp / *URA3* | This study |
| pCfB6912 | pCfB3042-URA; $P_{SNR52}$-gRNA(X-4)-$t_{SUP4}$ | Amp / *URA3* | This study |
| pCfB6915 | pCfB3045-URA; $P_{SNR52}$-gRNA(XI-3)-$t_{SUP4}$ | Amp / *URA3* | This study |
| pCfB6920 | pCfB3050-URA; $P_{SNR52}$-gRNA(XII-5)-$t_{SUP4}$ | Amp / *URA3* | This study |
| pCfB10292 | pCfB6910-URA; $P_{SNR52}$-gRNA(X-2, XI-3, XII-5)-$t_{SUP4}$ | Amp / *URA3* | This study |

**Table S2. Strains used in this study**

| Strain | Genotype | Marker | Parental strain | Reference |
|---|---|---|---|---|
| Sb | *URA3*Δ | N/A | SB-ATCC-796 | [2] |
| Sb GLP-1 (1x) | X-4 $P_{TDH3}$-GLP1-$t_{DIT1**}$ | N/A | Sb | This study |
| Sb GLP-1 | XI-3 $P_{TDH3}$-GLP1-$t_{DIT1**}$ | N/A | Sb | This study |

| | | | | |
|---|---|---|---|---|
| (2x) | XII-5 $P_{TDH3}$-GLP1-$t_{DIT1}$** | | | |
| Sb GLP-1 (4x) | X-2 $P_{TDH3}$-GLP1-$t_{DIT1}$**<br>X-4 $P_{TDH3}$-GLP1-$t_{DIT1}$**<br>XI-3 $P_{TDH3}$-GLP1-$t_{DIT1}$**<br>XII-5 $P_{TDH3}$-GLP1-$t_{DIT1}$** | N/A | Sb GLP-1 1x | This study |
| Sb Exe-4 (1x) | XI-3 $P_{TDH3}$-Exe4-$t_{DIT1}$** | N/A | Sb | This study |
| Sb Exe-4 (2x) | XI-3 $P_{TDH3}$-Exe4-$t_{DIT1}$**<br>XII-5 $P_{TDH3}$-Exe4-$t_{DIT1}$** | N/A | Sb Exe4 1x | This study |

**Table S3. Oligonucleotides used in this study**

| Oligo Name | Sequence | Reference(s) |
|---|---|---|
| Pepck_fw | GGCCACAGCTGCTGCAG | [3] |
| Pepck_rv | GGTCGCATGGCAAAGGG | |
| G6pc_fw | CTGTGAGACCGGACCAGGA | [3] |
| G6pc-rv | TAGTATACACCTGCTGC | |
| Ucp1_fw | AGGCTTCCAGTACCATTAGGT | [4] |
| Ucp1_rv | CTGAGTGAGGCAAAGCTGATTT | |
| Pgc1a_fw | CCCTGCCATTGTTAAGACC | [4] |
| Pgc1a_rv | TGCTGCTGTTCCTGTTTTC | |
| Tfam_fw | GTCCATAGGCACCGTATTGC | [4] |
| Tfam_rv | CCCATGCTGGAAAAACACTT | |
| Cd36_fw | TGTGTTTGGAGGCATTCTCA | [4] |
| Cd36_rv | TTTTGCACGTCAAAGATCCA | |
| Aco_fw | TGTTAAGAAGAGTGCCACCAT | [5] |
| Aco_rv | ATCCATCTCTTCATAACCAAATTT | |
| Ppara_fw | AGAGCCCCATCTGTCCTCTC | [6] |
| Ppara_rv | ACTGGTAGTCTGCAAAACCAAA | |
| Tbp_fw | TTCTCGAAAGAATTGCGCTGT | [6] |
| Tbp_rv | GCCTTGTGAGTCATTTCAGTGA | |
| Tfiib_fw | GTTCTGCTCCAACCTTTGCCT | [7] |
| Tfiib_rv | TGTGTAGCTGCCATCTGCACTT | |
| Hprt_fw | CAAACTTTGCTTTCCCTGGT | [8] |
| Hprt_rv | TCTGGCCTGTATCCAACACTTC | |

**Table S4. Construct sequences**

| Name | Sequence | Reference(s) |
|---|---|---|
| Upstream overhang | tctaccaacggaatgcgtgcgatcgcgtgcattc | This study |

| | | |
|---|---|---|
| $P_{TDH3}$ | cgagtttatcattatcaatactgccatttcaaagaatacgtaaataattaatagta gtgattttcctaactttatttagtcaaaaaattagccttttaattctgctgtaaccc gtacatgcccaaaatagggggcgggttacacagaatatataacatcgtaggt gtctgggtgaacagtttattcctggcatccactaaatataatggagcccgctttt taagctggcatccagaaaaaaaaagaatcccagcaccaaaatattgttttctt caccaaccatcagttcataggtccattctcttagcgcaactacagagaacag gggcacaaacaggcaaaaaacgggcacaacctcaatggagtgatgcaac ctgcctggagtaaatgatgacacaaggcaattgacccacgcatgtatctatct cattttcttacaccttctattaccttctgctctctctgatttggaaaaagctgaaaa aaaaggttgaaaccagttccctgaaattattcccctacttgactaataagtatat aaagacggtaggtattgattgtaattctgtaaatctatttcttaaacttcttaaatt ctacttttatagttagtcttttttttagttttaaaacaccaagaacttagtttcgaata aacacacataaacaaacaaa | This study |
| Kozak | aacaaa | [9] |
| Alpha-pre | atgagatttccatctatttttactgctgttttgtttgctgcttcttctgctttggct | [9] |
| Alpha-pro | gctccagttaatactactactgaagatgaaactgctcaaattccagctgaagc tgttattggttattctgatttggagggtgactttgatgttgctgttttgccattttcta actctactaacaacggtttgctattcatcaacactactatcgcttctatcgctgct aaagaagaaggtgtttctttggat | [9] |
| Kex2 site | aaaaga | [9] |
| Spacer | gaagaaggtgaaccaaaa | [9] |
| GLP-1 | catgacgaattcgaaagacacgcagaaggcacgtttacaagcgacgtgag ctcttaccttgaaggtcaagcagcgaaagaattcatagcatggttagtaaaag gtagaggctaa | This study |
| Exendin-4 | catggtgaaggcacattcacatctgatctgtccaaacaaatggaggaggaa gcggtacgtttatttattgaatggttaaaaaacgggggacctagctccggcgc gcccccccgagctaa | This study |
| $t_{DIT}$** | taaagtaagagcgctacattggtctaccttttctttttacttaaacattagttagtt cgttttctttttcttttttatgtttcccccccaaagttctgattttataatattttatttc acacaattccatttaacagaggggggaatagattctttagcttagaaaattagtg atcaatatatatttgcctttcttttcatcttttcagtgatattaatggtttcgagacac tgcaatggccct | [10] |
| Downstream overhang | actagtgctgaggcattaat | This study |
| Ordered gBlock GLP-1 | tctaccaacggaatgcgtgcgatcgcgtgcattccgagtttatcattatcaata ctgccatttcaaagaatacgtaaataattaatagtagtgattttcctaactttattt agtcaaaaaattagccttttaattctgctgtaacccgtacatgcccaaaatagg gggcgggttacacagaatatataacatcgtaggtgtctgggtgaacagtttat tcctggcatccactaaatataatggagcccgctttttaagctggcatccagaa aaaaaaagaatcccagcaccaaaatattgttttcttcaccaaccatcagttcat aggtccattctcttagcgcaactacagagaacaggggcacaaacaggcaa aaaacgggcacaacctcaatggagtgatgcaacctgcctggagtaaatgat gacacaaggcaattgacccacgcatgtatctatctcattttcttacaccttctatt | This study |

| | | |
|---|---|---|
| | accttctgctctctctgatttggaaaaagctgaaaaaaaaggttgaaaccagtt<br>ccctgaaattattcccctacttgactaataagtatataaagacggtaggtattga<br>ttgtaattctgtaaatctatttcttaaacttcttaaattctacttttatagttagtcttttt<br>tttagttttaaaacaccaagaacttagtttcgaataaacacacataaacaaaca<br>aaaacaaaatgagatttccatctatttttactgctgttttgtttgctgcttcttctgc<br>tttggctgctccagttaatactactactgaagatgaaactgctcaaattccagct<br>gaagctgttattggttattctgatttggagggtgactttgatgttgctgttttgcca<br>ttttctaactctactaacaacggtttgctattcatcaacactactatcgcttctatc<br>gctgctaaagaagaaggtgtttctttggataaaagagaagaaggtgaaccaa<br>aacatgacgaattcgaaagacacgcagaaggcacgtttacaagcgacgtg<br>agctcttaccttgaaggtcaagcagcgaaagaattcatagcatggttagtaaa<br>aggtagaggctaataaagtaagagcgctacattggtctaccttttcttttactt<br>aaacattagttagttcgttttctttttctttttttatgtttccccccaaagttctgattt<br>tataatattttatttcacacaattccatttaacagaggggggaatagattctttagct<br>tagaaaattagtgatcaatatatatttgcctttcttttcatcttttcagtgatattaat<br>ggtttcgagacactgcaatggccctactagtgctgaggcattaat | |
| Ordered gBlock Exendin-4 | tctaccaacggaatgcgtgcgatcgcgtgcattccgagtttatcattatcaata<br>ctgccatttcaaagaatacgtaaataattaatagtagtgattttcctaactttattt<br>agtcaaaaaattagccttttaattctgctgtaacccgtacatgcccaaaatagg<br>gggcgggttacacagaatatataacatcgtaggtgtctgggtgaacagtttat<br>tcctggcatccactaaatataatggagcccgctttttaagctggcatccagaa<br>aaaaaaagaatcccagcaccaaaatattgttttcttcaccaaccatcagttcat<br>aggtccattctcttagcgcaactacagagaacaggggcacaaacaggcaa<br>aaaacgggcacaacctcaatggagtgatgcaacctgcctggagtaaatgat<br>gacacaaggcaattgacccacgcatgtatctatctcattttcttacaccttctatt<br>accttctgctctctctgatttggaaaaagctgaaaaaaaaggttgaaaccagtt<br>ccctgaaattattcccctacttgactaataagtatataaagacggtaggtattga<br>ttgtaattctgtaaatctatttcttaaacttcttaaattctacttttatagttagtcttttt<br>tttagttttaaaacaccaagaacttagtttcgaataaacacacataaacaaaca<br>aaaacaaaatgagatttccatctatttttactgctgttttgtttgctgcttcttctgc<br>tttggctgctccagttaatactactactgaagatgaaactgctcaaattccagct<br>gaagctgttattggttattctgatttggagggtgactttgatgttgctgttttgcca<br>ttttctaactctactaacaacggtttgctattcatcaacactactatcgcttctatc<br>gctgctaaagaagaaggtgtttctttggataaaagagaagaaggtgaaccaa<br>aacatggtgaaggcacattcacatctgatctgtccaaacaaatggaggagg<br>aagcggtacgtttatttattgaatggttaaaaaacgggggacctagctccggc<br>gcgcccccccccgagctaataaagtaagagcgctacattggtctaccttttctt<br>ttacttaaacattagttagttcgttttctttttctttttttatgtttccccccaaagttc<br>tgattttataatattttatttcacacaattccatttaacagaggggggaatagattct<br>ttagcttagaaaattagtgatcaatatatatttgcctttcttttcatcttttcagtgat<br>attaatggtttcgagacactgcaatggccctactagtgctgaggcattaat | This study |

#### Supplementary methods

##### Real-time growth monitoring

Real-time $OD_{600}$ was measured every 10 min for approximately 48 h with microplate reader Synergy$^{TM}$ H1 BioTek. The cultures were incubated into a 200 µL DELFT medium supplemented with 20 mg/l uracil in a CELLSTAR® 96 well cell culture plate (Greiner Bio-One) with an air-penetrable lid (Breathe-Easy, Diversified Biotech). Cultivation was performed with continuous double orbital shaking of 548 cycles per minute (CPM) at 37°C and with an initial $OD_{600}$ of 0.05.

##### Animal studies

##### Characterisation of colonisation in conventional mice over 24 hours

The mice were divided into two groups, either receiving Sb-Empty (n = 4) or Sb-Exe4 (n = 4). All groups were orally administered via intragastric gavage daily in the morning for five days. Faeces were collected prior the first oral administration and on the fifth day at 3, 6, 9, 12 and 24 hours after the fifth oral administration.

##### *In vivo* characterisation of Sb-Exe4 in male C57BL/6 mice for 35 days

The mice were divided into two groups and were orally administered for 35 days with either Sb-Empty (n = 7) or Sb-Exe4 (n = 7). Body weight was monitored weekly. An intraperitoneal glucose tolerance test was performed on day 22. Faeces was collected on day 0, 7, 14, 21, 28, and 35. The mice were euthanised on day 35.

##### Synergetic effect of Sb-Exe4 and cold exposure on energy homeostasis

The mice were divided into two groups and were orally administered for 55 days with either Sb-Empty (n = 8) or Sb-Exe4 (n = 5 – 7). Body weight was monitored weekly. Faeces were collected on day 0, 7, 14, 21, and 28. An intraperitoneal glucose tolerance test was performed on day 22 and 44 (data not shown). The mice were moved into metabolic cages from day 29. On day 46 the mice were exposed to cold (8°C). The cold exposure was interrupted on day 49 to 52 due to a malfunction of the Climate Chamber. The mice were euthanised on day 55.

##### Glucose tolerance test (GTT)

Intraperitoneal glucose tolerance test (IP-GTT) was performed three weeks after the treatment for the antibiotic-treated mice and three and six weeks into the treatment for the conventional mice. Mice were fasted for 6 hours and injected intraperitoneally with a 20% glucose solution prepared in PBS (2 mg/g body weight). Blood was drawn from the tail vein before and after the glucose injection at 0, 15, 30, 60, 90, and 120 min. Blood glucose was measured using the Accu-Check Guide meter (Roche).

##### Western blotting

The sWAT were homogenised in a homogenisation buffer containing 50 mM HEPES, 1 % Triton X-100, 50 mM Na-pyrophosphate, 100mM sodium fluoride, 10 mM EDTA, and protease inhibitor cocktail (Sigma Aldrich Cat. #P8340). Supernatant was collected and quantified with Bradford Protein Quantification (Bio-Rad DCTM Protein assay). Four random samples were selected for further analysis. Equal amounts of proteins were separated by 10% SDS-PAGE and blotted onto a polyvinyl difluoride (PVDF) membrane. The membranes were blocked in 5 % skim milk in 1 % PBS for 60 min, probed with primary antibody at 4°C overnight. The membrane was washed three times (3x 10 min) in TBST before incubated with secondary antibody for 60 min. The membranes were washed in TBST for three times (3x 10 min) before ECL detection. The antibodies used were anti-UCP1 (1:5000) (Abcam, Cat. #ab209483), anti-ß-Tubulin (1:500) (Abcam, Cat. #ab6046), anti-rabbit IgG-HRP (1:3000) (Santa Cruz, Cat. #sc-2357-CM) and m-IgGκ BP-HRP (1:3000) (Santa Cruz Cat. #sc-516102-CM).

**Histology**

Adipose tissues (iBAT and sWAT) were dissected, fixed in 4% formaldehyde, embedded, and sectioned into 5 mm slides. Three to four random samples were selected for further analysis. Sections were de-waxed in xylene and hydrated in a series of decreasing ethanol concentrations. Afterwards, antigen was retrieved by immersing the slides in 10 mM sodium citrate (pH 6.0) buffer and heating in the microwave (300W) for 20 min. Slides were then rinsed in water and immunostained for UCP1 using a Vectastain ABC kit (Vector Lab, Cat. #PK-6101, rabbit IgG) following the protocol provided by the manufacturer. Briefly, the slides were first blocked in the PBS/serum blocking buffer (ABC kit), then incubated with Anti-UCP1 antibody (Abcam, Cat. #ab209483, 1:200 dilution) at 4°C overnight. The next day, slides were washed with PBS and incubated with the anti-rabbit 2$^{nd}$ antibody (from the ABC kit), followed by incubation with the ABC reagent (from the ABC kit). Eventually, colour staining was developed by incubating the slides in a prepared DAB substrate reagent (Vector Lab, Cat. #SK-4100). Finally, the slides were counterstained with haematoxylin, washed in water, and mounted in an antifade mounting medium. Adipocyte cell number and size were analysed by ImageJ.
